# Best practices for cryo-trapping time-resolved crystallography with the Spitrobot crystal plunger

**DOI:** 10.64898/2026.04.07.716871

**Authors:** Robert Bosman, Caitlin E. Hatton, Andreas Prester, Maria Spilliopolou, Friedjof Tellkamp, Pedram Mehrabi, Eike C. Schulz

**Author notes:** Corresponding author April, 2026. These authors contributed equally (Robert Bosman, Caitlin E. Hatton, Andreas Prester).

## Abstract

Capturing meta-stable conformations of enzymes and ligand complexes demands structural snapshots beyond static crystal structures. While time-resolved serial crystallography at room temperature, offers a time-resolution down to the femto-second domain it requires large amounts of micro crystals, specialized beamlines and considerable experience. Moreover, as the majority of enzymes displays turnover-times in the millisecond domain or slower, simpler methods can provide meaningful structural insight into enzyme catalysis. Vitrification of protein crystals can trap reaction intermediates by rapid cooling to ¡ 100 K, and has traditionally been used to gain insight into long lived reaction intermediates such as product complexes. However, manual vitrification procedures are limited to long delay times of at least several seconds and heavily suffer from operator variability. A solution to this problem is provided by automatic crystal plunging devices, such as the Spitrobot, that plunge loop-mounted protein crystals into liquid nitrogen within millisecond time-scales. Versatile means of reaction initiation can be achieved either by micro dispensing a ligand droplet, or via optical excitation of light-sensitive proteins, or via the photoactivation of caged compounds. In addition to the conceptual simplicity, another benefit of cryo-trapping is that data can be collected at conventional synchrotron beamlines, exploiting their robust high-throughput capabilities. Thus, compared to room-temperature time-resolved crystallography, users not only benefit from uncoupling sample-preparation and data-collection, but also from a reduction in the required technical expertise and ready access to radiation sources. However, as cryo-trapping crystallography explores dynamic structural changes that become only visible by the comparison of several samples, experiments have to be carefully planned to carry out the necessary controls and to avoid mis- or over-interpretation of the results. Here we describe a detailed protocol for cryo-trapping time-resolved crystallography using automated crystal-plungers that enables researchers to map enzymatic reaction coordinate pathways within the millisecond domain.

## 1 Introduction

To understand the molecular foundations of life, it is imperative to understand the protein dynamics which underpin various biochemical functions. High-resolution protein structures of meta-stable functional states provide important information on, ligand binding, cooperative mechanisms, allosteric regulation, detailed catalytic chemistry, and most crucially the interrelations between these phenomena. To understand enzyme chemistry this detail is required to fully interpret the reaction mechanism and the dynamic structural changes required to permit catalysis. Cryo-trapping of intermediate species *in-crystallo* is a powerful, yet conceptually simple technique in structural biology for answering these questions.

X-ray crystallography provides protein structures at high spatial precision but these are typically static or at steady state. To overcome this, protein function can be initiated *in-crystallo*, by e.g. photoactivation [1, 2], or ligand diffusion [3–12]; alternatively, perturbations such as pH jumps [13] or release of solvated electrons [14] can be utilised [15]. Cryo-trapping then stops these reactions by rapidly cooling the sample to below the glass transition phase (∼100 K) [16–18]. This works serendipitously with the common practise of cryo-cooling samples to decrease radiation damage and increase crystal-life times in the beam.

Cryo-trapping thus enables structural characterization of meta-stable states along enzymatic or photochemical reaction pathways. Historically, it has provided essential insights into a wide range of systems [19, 20]. Particularly, interrogating slow enzymes with turnover durations from minutes to seconds [21, 22], extending our knowledge from apo, product, or dead-end complexes, to functional intermediates only present under out-of-equilibrium conditions.

Recently, serial crystallography, where partial diffraction patterns from thousands of micro-crystals are merged, has allowed data-collection with minimal radiation damage at non-cryogenic temperatures [23, 24]. This has been expanded to time-resolved studies with each crystal initiated separately, merged serial datasets collected over multiple time-delays construct a series of structures along a reaction coordinate. Soon after this strategy was successfully applied at X-ray free-electron lasers (XFELs) enabling serial-femtosecond crystallography, it was also successfully adapted at synchrotron radiation sources for time-resolved analyses of enzymatic reaction mechanisms [25–32].

The adaptation to synchrotron beamlines was made possible by the development of microfocus beams and improved detectors, allowing usable signal from smaller crystals [33]. Time-resolved crystallography in-general, benefits from a reduced crystal size, which improves the homogeneity of reaction initiation. In photo-triggered reactions, the crystal is more uniformly exposed to photons by reducing the absorption depth [34, 35], while in ligand mixing experiments it shortens the diffusion time to the crystal centre and thereby improves the stoichiometry with the ligand [4, 36–39]. For cryo-trapping techniques, this reduced crystal size results in a more rapid equilibration than previously possible, enabling cryo-trapping at time-scales more akin to RT time-resolved crystallography. Time-resolved cryo-trapping experiments main advantages lie in the general simplicity, wide applicability, and compatibility with high-throughput data-collection. Rapid diffusion cryo-trapping experiments have wide potential in structural biology, biochemistry, and mechanistic enzymology. Alternatively they provide new opportunities for fragment screening, inhibitor design, and early-stage drug discovery. The simplest version of the method requires only standard crystallographic tools, any MX laboratory can adopt the basic manual workflow.

Unique to cryo-trapping experiments is, that the crystals size is also relevant to the vitrification time. Microcrystals in the range of 10-50 *µ*m are typically optimal to minimise vitrification times, improving the overall homogeneity of reaction intermediates [40–43]. Additionally, studies have demonstrated that significant heat removal occurs in the cold gas layer above the cryogen, and plunge speeds of ∼1 m s*^−^*^1^ across a 2 cm gas gap can dominate the cooling process [44, 45] – further complicating manual trapping experiments. Eliminating or minimizing this gas layer via automated approaches, thus significantly enhances vitrification speeds [46]. These consideration are unique to cryo-trapping experiments making the time resolution and quality of trapping dependant on crystal size, sample environment, and plunge-freezing conditions. However, recently developed high-speed crystal plungers address these concerns, which enables trapping of enzymatic reaction intermediates in the millisecond time domain[43, 47–50].

Commonly, RT and cryo-trapping time-resolved crystallography aim to determine protein conformational states and intermediates. Whether intermediates can be observed is fundamentally contingent on whether a non-equilibrium state can be sufficiently populated *in-crystallo*. This depends on the crystal form, an appropriate initiation strategy, the intermediate thermodynamics, and data collection strategy. Automated plungers such as the *Spitrobot* do not change these underlying biochemical or physicochemical determinants, but facilitate measurements at time-scales not accessible to manual cryo-trapping. Additionally they simplify and improve the sample handling, and improve the measurement reliability. Given the growing adoption of time-resolved crystallography it is important to delineate how these results relate to cryo-trapping results.

Cryo-trapping has two distinctions to RT serial approaches. Firstly, it permits a rotation data collection strategy via increased radiation tolerance thus improving structure factor approximation with current methods. Secondly, the rapid cooling can shift the protein conformational landscape towards lower entropy states, which can induce cryo-cooling artifacts and increase sample heterogeneity due to the difference in freezing times [42]. As such cryo-trapping provides better data quality for more thermodynamically stable states, while serial approaches can capture higher-energy, meta-stable states that maybe quenched during freezing. This makes cryo-trapping highly complementary to time-resolved serial crystallography measurements [18, 51]. Therefore, the automated cryo-trapping, provided by automated plungers such as the *Spitrobot*, improves the experimental methodology and workflow simplicity. This makes it a useful initial measurement, before attempting more complicated serial experiments.

Here we summarize practical guidelines and protocols for sample preparation and data collection in cryo-trapping time-resolved crystallography. We initially focus on general considerations and important aspects in conventional (i.e. manual) cryo-trapping. Later we focus our description on cryo-trapping experiments using mechanical high-speed crystal plungers, highlighting our recently introduced *Spitrobot* as an example. The *Spitrobot* is targeted at equipping biochemistry facilities with an automated crystal-plunger platform to permit substantially shorter delay times and improved reproducibility, enabling mapping of enzymatic and ligand-binding kinetics across millisecond time-delays. The protocol stands to expand the user base for time-resolved structural biology and enhance the mechanistic insights achievable in structure-guided discovery pipelines. The workflow is applicable to mesophilic and thermophilic enzymes, multi-domain proteins, and complexes of biomedical, industrial, or biotechnological relevance. Reaction-initiation strategies include ligand delivery via droplet deposition or optical excitation of light-sensitive systems. The protocol incorporates environmental control of temperature and relative humidity, enabling users to maintain crystal stability across diverse protein classes while fine-tuning solvent content and lattice integrity.

## 2 Preliminary Experiments

Several experimental choices and optimisations need to be addressed before proceeding to more complicated cryo-trapping experiments, either as rapid manual or automated cryo-trapping measurements. An overview and order of these considerations is provided in Figure 1.

**Figure 1:**
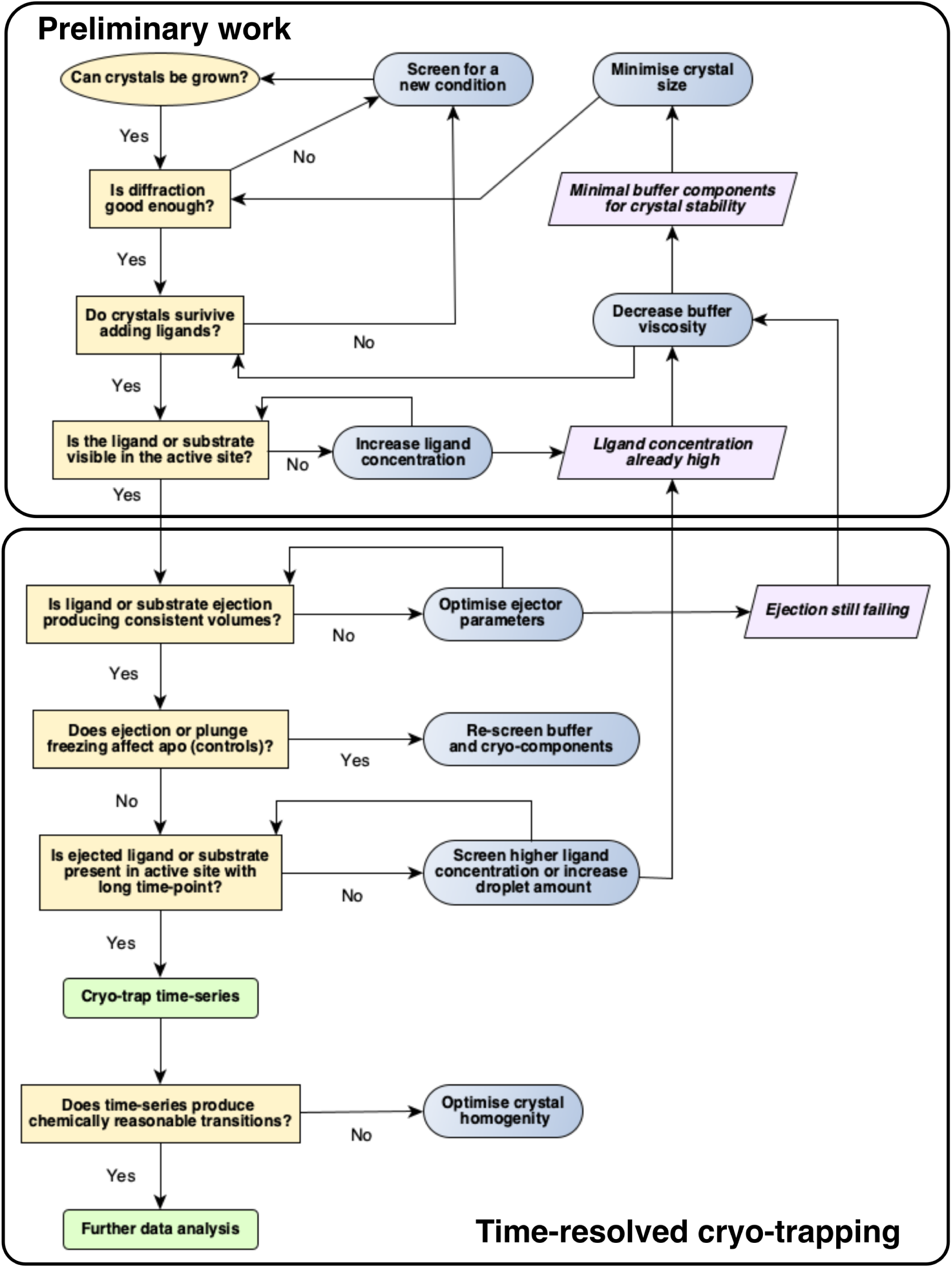
An overview of the experimental workflow. The work is split into two separate phases, the pre-experimental work, that includes sample optimisation and confirmation of ligand binding, and the time-resolved crystallography where a time-series of structures is feasible under common conditions.

### 2.1 Crystal growth and diffraction properties

As a central aim of the workflow is the comparison of multiple states at different delay-times after reaction initiation, crystal growth has to be reliable and lead to comparable, high quality diffraction patterns. Depending on the nature of the change, as a rule-of-thumb the resolution should not be lower than 2.5 Å. Obviously any crystallographic pathologies like e.g. twinning heavily complicate the downstream data-analysis and should be avoided.

While the initial manual experiments can be carried out using macroscopic crystals, its imperative that time-resolved experiments are carried out on crystals that permit homogenous reaction initiation typically these have micrometer dimensions. Protocols for homogenous micro-crystallization are identical to those required for RT-TRX and have been previously addressed in focussed publications [52–56]. These micro-crystallization conditions should be carefully optimized to achieve the best trade-off in diffraction properties and reaction initiation, as these are fundamental limits to all downstream experiments Figure 2a.

**Figure 2:**
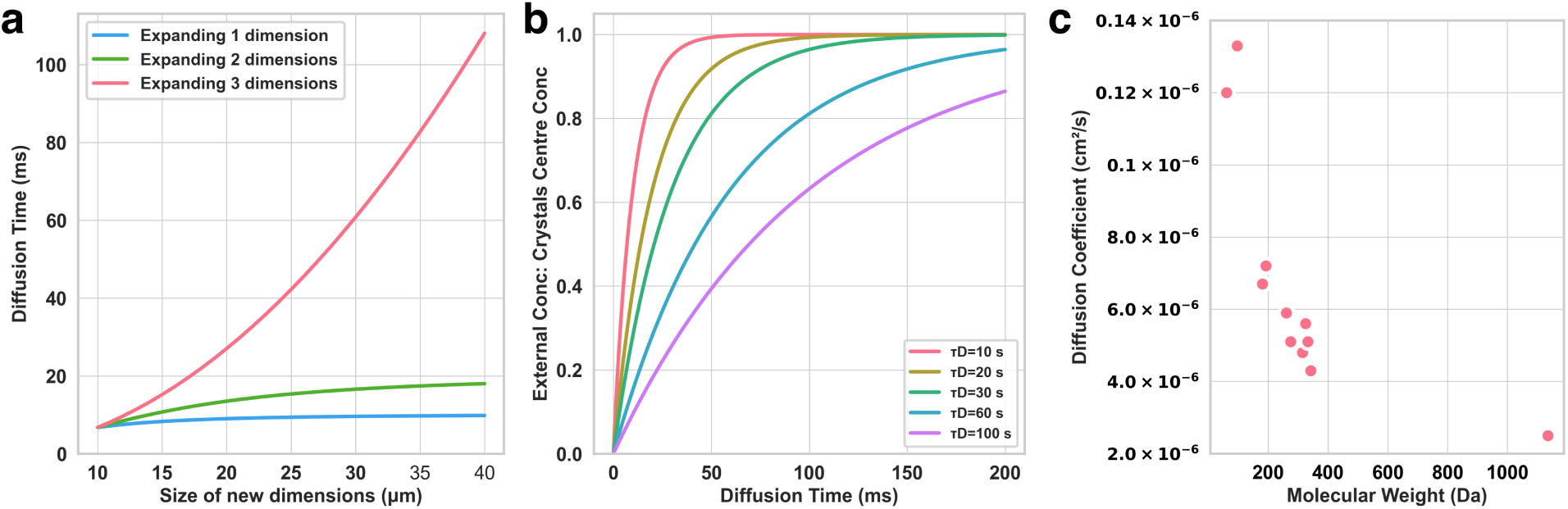
Diffusion considerations, calculations are adapted from [36], a) graphs the effect of expanding crystals dimensions on the diffusion time (*τ_D_*) as defined in [36], with *D* = 6 × 10*^−^*^6^. Note this graph highlights that rod or plates shaped crystals can have fairly long dimensions while diffusion is still relatively fast. b) The ratio between the external substrate concentration and concentration at the crystal centre for different diffusion times. b) It is often desireable to use a high substrate concentration to reduce the difussion time it takes to reach a given concentration. As the graph indicates, if the desired concentration to achieve an good stoichometry is 20% of the ligand concentration in the droplet, even with a long overall diffusion time (120 ms) it would take ∼25 ms to reach the desired concentration. Therefore applying high ligand concentrations can in-part mitigate slow difussion kinetics. c) The coefficients for several small molecules plotted agaisnt there molecular weight (data from [**a**, **b**]). This plot highight the correlation between molecular weight and faster difussion coefficient and therefore a smaller molecule is generally preferable. There are caveats to this, such as charged ions, and we point the reader to more rigorous calculations and a dicussion on diffusion into crystals provided in[58]

### 2.2 Cryo-protection

Unlike room-temperature time-resolved crystallography, cryo-trapping experiments typically require the addition of cryo-protectants to prevent the formation of ice crystals and damage to the crystals during freezing [57]. These are typically organic solvents, sugars, or different molecular weight PEGs, all of which increase buffer viscosity. This poses a challenge for both optimal ligand diffusion and droplet injection. Therefore, cryo-protectants should be optimised to reduce the concentration and utilize less viscous cryo-protectants such as organic solvents or low-molecular weight PEGs.

### 2.3 Soaking experiments

Once apo protein crystals can be grown and give sufficient data-quality with an optimised cryo-protectant, manual soaking experiments are essential to confirm substrate binding. These follow standard practices in crystallography, with ligands added directly to the crystallization well or fished into droplets with cryo-protectant and ligand prior to flash-freezing. A key reason for utilizing faster more precise cryo-trapping methodologies is that enzymatic turnover is often faster than can be trapped manually. Both rapid turn-over and an occluded active site produce an empty active site. In this instance biochemically assaying the protein crystals can confirm catalysis within the lattice [4]. Reducing the crystal size, decreases the equilibration times Figure 2a, alternatively soaking at lower temperatures can slow the reaction enough to confirm binding and turnover [4, 5]. Ensuring that a product state is observed via manual soaks acts as an essential positive control for ligand mixing experiments. If no evidence of activity in crystals is viable, then re-screening for a new crystal form is required.

### 2.4 Substrate and Ligand considerations

When considering ligand addition reactions there is an additional choice of initiating ligand or substrate. As the *Spitrobot* enables mixing times which would be manually unfeasible, it permits working with native substrates that catalyse too quickly for manual cryo-trapping. This minimizes the need to work with mutants which often lack key chemical groups at important residues that would otherwise predominate the native pathway. Despite this, diffusion is often a limiting factor within the confines of the crystal lattice. When considering potential substrate or ligands there are three preferred characteristics: high solubility, high diffusion coefficient, and a poor substrate as defined by *k_cat_*/*K_M_*(in-order for catalysis to be slower than diffusion) . This results in preferential ligand properties to consider when designing a mixing experiment Figure 2.

### 2.5 Delay-times

Assuming that an observable product-state has already been achieved via manual soaking, different strategies can be used for selecting time-delays. If *in-crystallo* kinetics are known, (e.g. assessed spectroscopically) it is straightforward to start with suitably populated time-delays. If *in-crystallo* kinetics are unknown, time-delays can be decreased from the initial control following a quasi-logarithmic spacing to a few-hundred milliseconds [29]. Initial data-analysis should indicate when events are occurring *in-crystallo* and further time-delays can be planned accordingly.

## 3 Control Experiments

Before moving from manual to automated cryo-trapping, several control measurements are important to ensure that changes can be correctly attributed to the added ligand (Figure 3). These controls are focused around ensuring the automated cryo-trapping process is not interfering with the samples. In contrast to canonical single-crystal structure determination experiments, it is vital to carry out these controls for cryo-trapping experiments. For a meaningful evaluation of the data, it has to be verified that any observed electron density changes originate from a biochemical function and are not simply experimental artefacts due to the perturbation of the crystals. These controls adequately apply to any automated plunge freezing device and are recommended initial measurements on the *Spitrobot*.

**Figure 3:**
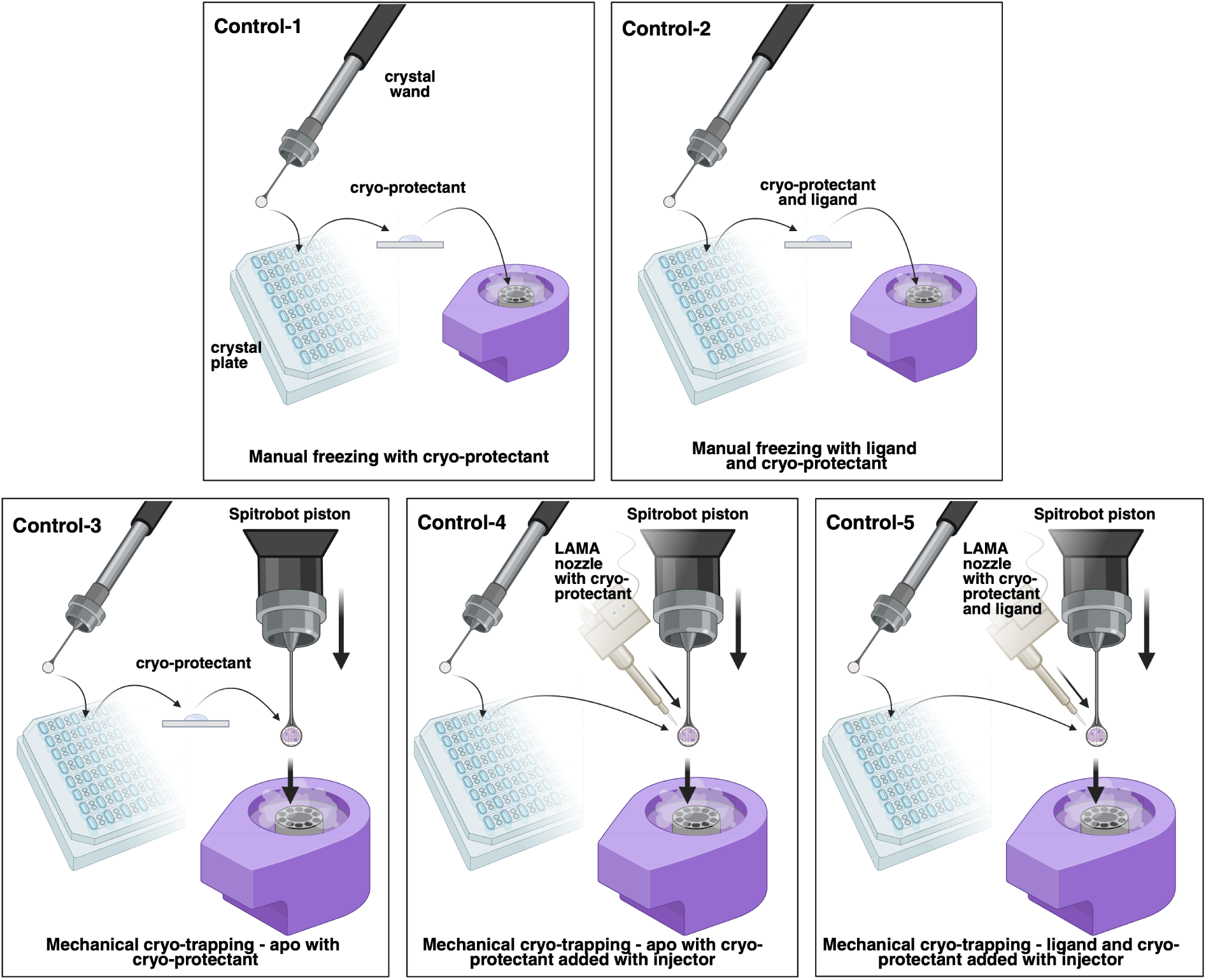
Schematic overview of the controls. Controls-1, manually freezing the sample with cryo-protection should ensure that the cryo-protectant is sufficient as usual in crystal screening, but uniquely whether the cryo-protectant binds or may intefere with the time-resolved study. Control-2, is a soak with the substrate. If the substrate is not present re-screening is likely required. Control-3 the sample can be mounted normally, but spitting is excluded. If the plunge freezing is causing an adverse effects it should be visible here. Control-4 the spitter is used to add cryo-protectant rather than a soak before mounting. Idealy Control-1, 3, and 4 will produce identical maps. Finally, Control-5 is essentially a long time-point measurement. This should produce similar results to manual control-2.

### Control-1: Manually frozen with cryo-protectant

Collecting a unperturbed apo state without change to the sample, optimized cryo-protectant is the initial baseline measurement. Establishing a coherent baseline can be achieved by measuring multiple crystals to similar data quality. This point is also ideally suitable for the selection of an appropriate crystals mount, discussed in further detail in section 8.1.

### Control-2: Manually frozen with ligand or substrate and cryo-protectant

Ligand or substrate can be added and cryo-trapped manually. The most straightforward approach is transferring the crystal into a well containing the ligand and freezing in liquid nitrogen after several seconds. This is essentially the soaking experiment that is a key part of the preliminary work, we should see a product state or evidence of binding and catalysis in the active site. Please refer to section 5.1 for a detailed outline of the procedure.

### Control-3: Mechanical cryo-trapping - Apo samples with cryo-protectant

Prior work details that the method of freezing can change the characteristics of the sample [59]. Therefore, when starting an automated cryo-trapping approach, controls should be measured to ensure that utilizing the device recapitulates the data quality and electron density from original apo states. Crystal damage due to insufficient humidity, temperature control or the rapid plunge freezing process should be detected from this control. Additionally, the higher vitrification speed provided by automated plungers may reduce the required cryo-protectant concentration.

### Control-4: Mechanical cryo-trapping - Apo samples with cryo-protectant added via systems liquid application method (LAMA nozzle)

Other sources of error can come about from distortions caused by adding the cryo-protectant with the ligand via droplet addition Figure 7, particularly if the cryo-protectant concentrations differ. Therefore control measurements are required to characterise any signal artifacts caused by cryo-protectant addition rather than the ligand.

### Control-5: Mechanical cryo-trapping - Ligand and cryo-addition via droplet addition

This series of controls will be able to confirm that interpreted density is caused purely by ligand addition. Therefore the final control is to re-capitulate the ligand complex observed with the manual soaks established before proceeding to automated cryo-trapping using the plunge freezing device. An important consideration here is that the ligand stoichiometric ratios are in similar regime to ensure comparable maps.

## 4 Time-resolved cryo-trapping with the *Spitrobot*

The previous sections addressed a series of general considerations and required preparatory work before attempting time-resolved cryo-trapping experiments. We will now give a brief introduction to the *Spitrobot* system [48–50] , which we use as an example for automated cryo-trapping devices used in this protocol.

The *Spitrobot* utilises the LAMA method to enable accurate and reproducible cryo-trapping experiments with versatile time resolutions and the use of SPINE standard pins for compatibility with most synchrotrons [28]. It is equipped with a humidity flow device (HFD), which supplies a constant air flow at a preset temperature and humidity, allowing for multi-temperature crystallography and a suitable protein crystal environment. The effect of structural changes during flash-cooling [42] is reduced due to the rapid and repeatable plunge-freezing of the *Spitrobot* compared to manual flash freezing [50].

The *Spitrobot* setup comprises three principal parts. The *Spitrobot-2* device, the ancillary devices, and a driving computer. The main device as outlined in [50] internally integrates the following principal components; i) a standard-pin mount on a motorised piston, ii) a mounting stage with XYZ motion for the Autodrop pipette, iii) a sample shutter system, iv) a LN_2_ dewar mount, v) a puck rotation and alignment system, vi) two camera’s with XY alignment stages for sample visualisation and validation images. This devices also houses the electronics for coordinating the devices during automated cryo-trapping. The device is principally controlled by three buttons located on the front panel B1 and B2 to trigger the device. The third button, B3, is designed to cycle between a sample visualisation mode by providing diffuse background light, puck alignment mode by switching a guide beam, and neither state. It is also pressed to reset the spitrobot from its error state (Figure 6c)

The two main ancillary devices are the HFD, and a triggering system, principally the Autodrop pipette (LAMA nozzle) from microdrop technologies for which an in-built mount is incorporated (Figure 4b,c). Prior to performing experiments the ancillary devices need to be setup and mounted to the *Spitrobot-2*. The Autodrop pipette ejection parameters need to be re-optimised for a new buffer, as drop-ejection parameters change with the buffers physiochemical properties, we outline a specific Autodrop ejection optimisation protocol (SI Figure 1). Separately, the HFD needs to be equilibrated to the desired temperature and humidity approximately 30 mins before running the *Spitrobot*. Recent work has incorporated an optical trigger system [49] currently position via an custom mount to where the ejector sits. Finally, a control laptop is used to set the experimental parameters and monitor the LN2 level. Time-delays for camera triggering, pin-mount position height, sample numbering, validation image output folders, and sample time-delay are all specified via a LabView interface [48] from the laptop.

**Figure 4:**
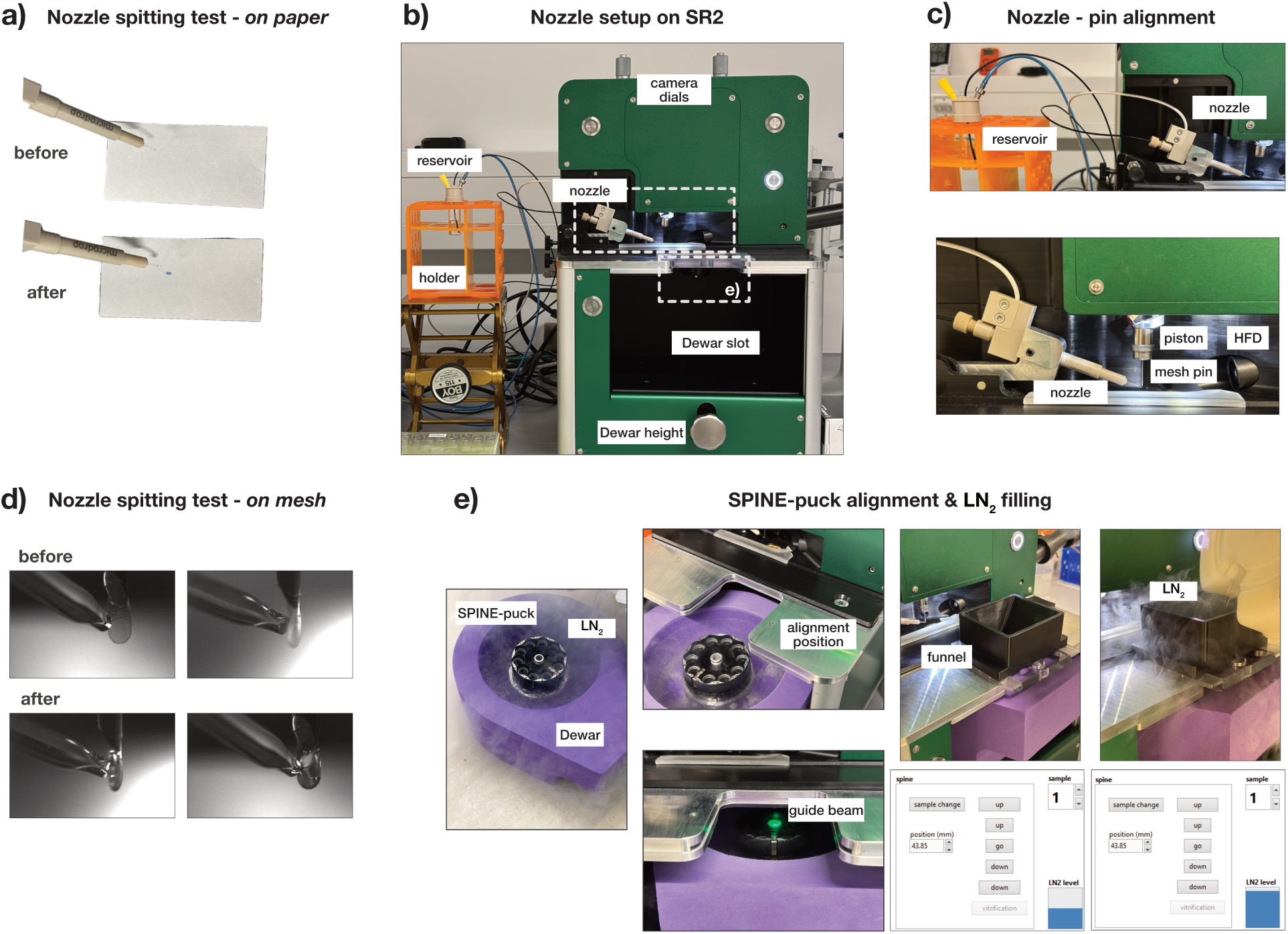
*Spitrobot* setup, a) Before and after successful ejection from the Autodrop pipette assessed using Wator paper. This water sensitive paper turns into a strong blue on contact with water and therefore gives a clear indication of successful ejection . The nozzle can then be mounted on the *Spitrobot-2* ; b) the nozzle points into the sample position from the left, while the HFD nozzle is postioned orthogonally. In order to maintain an appropriate pressure in the tubing the buffer resevior is mounted on a jack that can be adjusted accordingly (Table 1) . The sample piston positions the sample vertically between both c). Once the Autodrop pipette and the sample mesh are aligned on camera d) different ejection volumes can be tested looking for is a thin film of ligand sample that covers the mesh. e) With the HFD and Autodrop mounted and aligned the SPINE puck can be cooled and mounted into the *Spitrobot*, a step-by-step guide is provided in the protocol section.

**Table 1:**
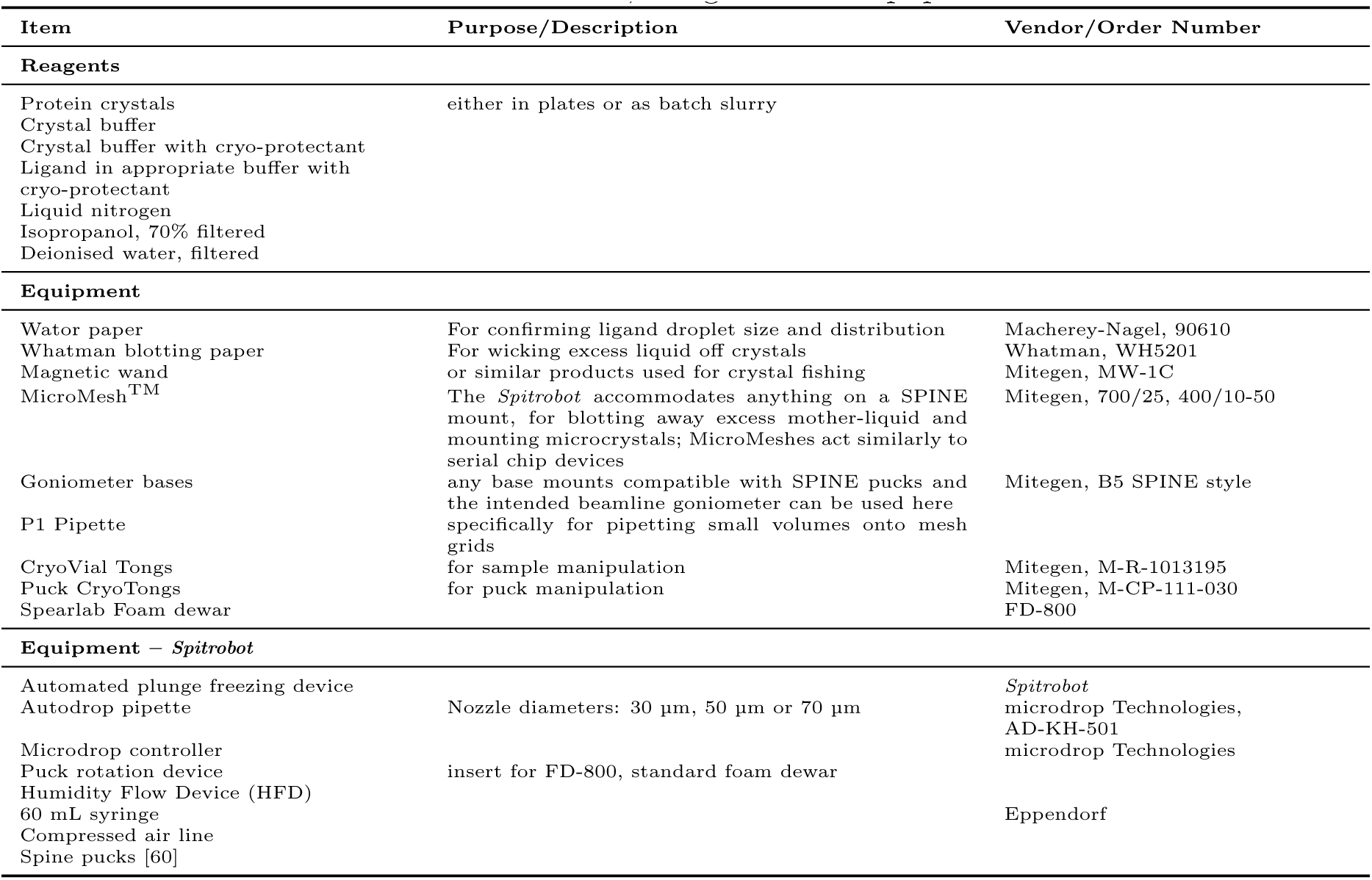
Materials, Reagents and Equipment.

## 5 Protocols & Procedures

### 5.1 Manual Cryo-trapping protocol

There are multiple versions of manually adding ligand and rapidly plunge cooling crystals. Here we outline a simple version for a microcrystal slurry that mimics the process achieved on the *Spitrobot* and therefore works well for controls 1 and 2. We would expect that a 10 s soaking time would be achievable. If product or end-states are already well explored crystallographically then skip to setting up ancillary devices.

1. Prepare the liquid nitrogen foam dewar and a puck as usual.

2. Mount the micromesh onto a magnetic wand, place the Whatman paper on the surface below the underside of the micromesh.

3. Pipette 0.25-1 µL microcrystal slurry onto the frontside of the mesh, aim to achieve a visible meniscus.

4. Refill the pipette with 0.25-1 µL containing ligand or buffer solution and hold the pipette tip just above the frontside of the mesh and tracking with the mesh in the next steps

5. Gently pivot the magnetic wand on the rim of the base, until the underside of the mesh contacts the whatman paper and absorbs the majority of the liquid. Raise the mesh just off the blot paper.

6. Pipette the ligand or buffer solution and repeat step 4.

7. Immediately, or after the pre-requisite time plunge the mesh into liquid nitrogen.

### 5.2 *Spitrobot* setup and configuration

Once all the ancillary devices are setup and mounted the *Spitrobot-2* device can be setup.

#### 5.2.1 Setting up and Mounting SPINE puck

8. **Critical**: Ensure the SpearLab foam dewar and puck rotation system are completely dry. Residual liquid left in the puck rotation system will freeze the mechanism. Other residual liquid will create floating ice crystals that can adhere to samples and compromise data collection.

9. Place the puck into holder and fill with liquid nitrogen. Once the puck has stopped boiling away liquid nitrogen, using the cryovial tongs rotate the puck manually to check the mechanism is still operational.

10. Align the puck to position the angular positioning groove faces towards the foam dewar handle. Place the foam dewar onto the inbuilt labjack.

11. Press the B3 button until the puck alignment light turns on (a green light that illuminates the centre position for the puck). Align the puck centre to the green light (Figure 6e).

12. Rotate the sample dial until in sample position ’7’. Align the angular positioning groove with a corresponding spoke located on the underside of the heating plate. Raise the labjack until both the puck centre and sample dial spoke are connected.

#### 5.2.2 Starting the *Spitrobot*

13. Ensuring the *Spitrobot* is powered on, start the control software. Once the *Spitrobot* has finished its internal checks. Request a shutter ’open’, and shutter ’closed’ to ensure the *Spitrobot* is properly connected.

14. **Recommended** Open the shutter and check the puck position matches the sample dial, rotate the puck to confirm the puck mechanism is rotating the puck (Figure 6c)

15. Rotate the puck to sample position 1.

16. If the Autodrop is not already mounted, mount the Autodrop pipette in the holder opposite the HFD and reconfirm sample spitting. Move the Autodrop pipette to the retracted position for more space around the piston.

### 5.3 Cryo-trapping samples with the *Spitrobot*

We outline the core procedure for automated plunge freezing using the *Spitrobot*, we assume that the spitrobot is already setup and configured. The HFD should be equilibrated to the set temperature and humidity, the SPINE puck should be loaded into the *Spitrobot* and set to position 1, with the Autodrop pipette already mounted.

### Recommended

Test plunging and spitting on an empty loop, try and get even coverage of ligand. (Figure 4d). For each solution it is important to optimize the amount of substrate deposited as excess ligand solution will contribute to background.

#### 5.3.1 Requesting the cryo-trapping parameters

17. In the LabView menu request the time-delay in milliseconds.

18. Using the dials align the nozzle in the X, Y, Z direction.

19. Set the pre-camera timings, as the number of milliseconds before plunging the cameras should image. For long time-points this can remain 200 ms, at timepoints ¡ 200 ms a minimum of 25 ms before plunge is recommended.

#### 5.3.2 Sample mounting and Alignment

20. There are two principal ways to mount samples depending on whether they are prepared in batch or are grown in a plate. The mounting procedure has small but important differences.

A. Microcrystals grown in batch

i. Mount the micromesh to the *Spitrobot* piston.
ii. Align the cameras so the micromesh is visible on the left-side of the image and in focus
iii. Rotate the mesh so the concave side faces the Autodrop pipette
iv. Pipette 0.25-1 µL of microcrystal slurry onto the front of the mesh.
v. Blot the convex mesh side with Whatman paper until only residual liquid remains.
B. Crystals fished into loops from plates

i. Fish the crystal from the wells and quickly mount to the *Spitrobot* piston to minimise drying out.
ii. Align the cameras so the loop is visible on the left-side of the image and in-focus
iii. Rotate the loop so that it is face onto the Autodrop
iv. Carefully blot-crucially without touching the crystal as much mother liquor as possible.

21. Detach the Autodrop pipette from its retracted position and move forward onto the right-side of the camera view. Use the Autodrop pipette XYZ stage to position the pipette tip above pointing down towards the centre of the sample mount (Figure 4c, d).

22. Align the Autodrop pipette to be inline with the sample, looking down the length of the pipette adjust pipette tip. The pipette tip should also come into focus of the camera.

#### 5.3.3 Triggering the *Spitrobot*

23. Depress the two trigger buttons located on the front panel of the *Spitrobot* with both hands to initiate plunge freezing. If plunge freezing needs to be halted release the trigger buttons. The buttons on the front panel will turn red to indicate the process has been halted (Figure 6b).

- **CAUTION** The freeze plunging mechanism is driven by an electric motor with a piston that is capable of piercing skin with the sample pin. The trigger buttons are designed be pushed with both hands to prevent the experimenters hands getting in the way of the mechanism while plunging. Others assisting with sample preparation should also stand back from the device while plunging.

24. ***Spitrobot* automated rapidly cooling process:** The automated vitrification takes place as outlined in previous publications [48, 50]. In brief the cameras take an image of the sample a set time before liquid deposition. The Autodrop pipette will deposit ligand. The camera will take another image to confirm deposition. If the time-point is short the shutter will open prior to these steps, and the sample will be immediately plunged, for longer time-points the shutter will open after deposition. The sample will remain in the down plunged position for several seconds before the magnet releases the sample into the puck. The plunger will then return to its home position and the shutter will close. Several issues can occur during this process, as outlined in Figure 5, a full guide on how to handle these can be found in the Troubleshooting table (Table 1).

**Figure 5:**
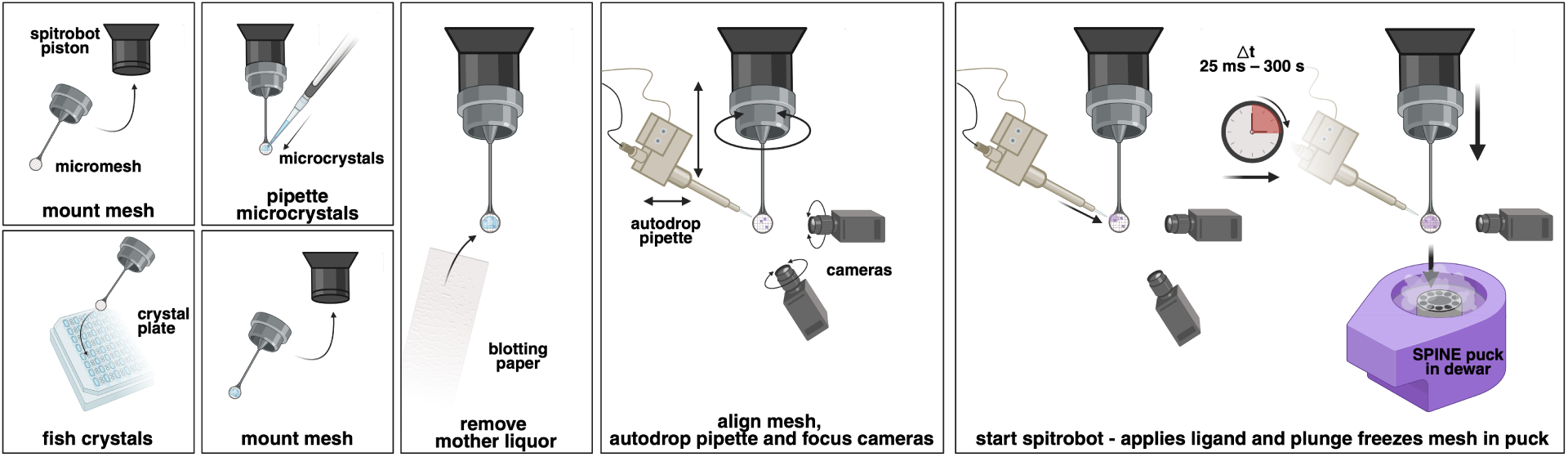
Schematic overview of the automated cryo-trapping workflow. Crystals can be pipetted directly onto a MicroMesh from a slurry, or fished from a crystallisation tray. Excess mother liquor should then be removed by blotting from the back side of the mesh. The mesh face with the deposited crystals is positioned in front of the Autodrop pipette, which is aligned closely with it. The spitrobot then automatically triggers the Autodrop pipette to apply ligand solution and vitrifies the sample in liquid nitrogen at the set time-delay.

25. Confirm droplet deposition via images taken before and after Autodrop triggering, before continuing to the next sample.

#### 5.3.4 Resetting the *Spitrobot*

26. Rotate the puck dial to the next sample position. It is recommended to manually open and close the shutter to confirm the mechanism has rotated for the first few samples (Figure 6c).

**Figure 6:**
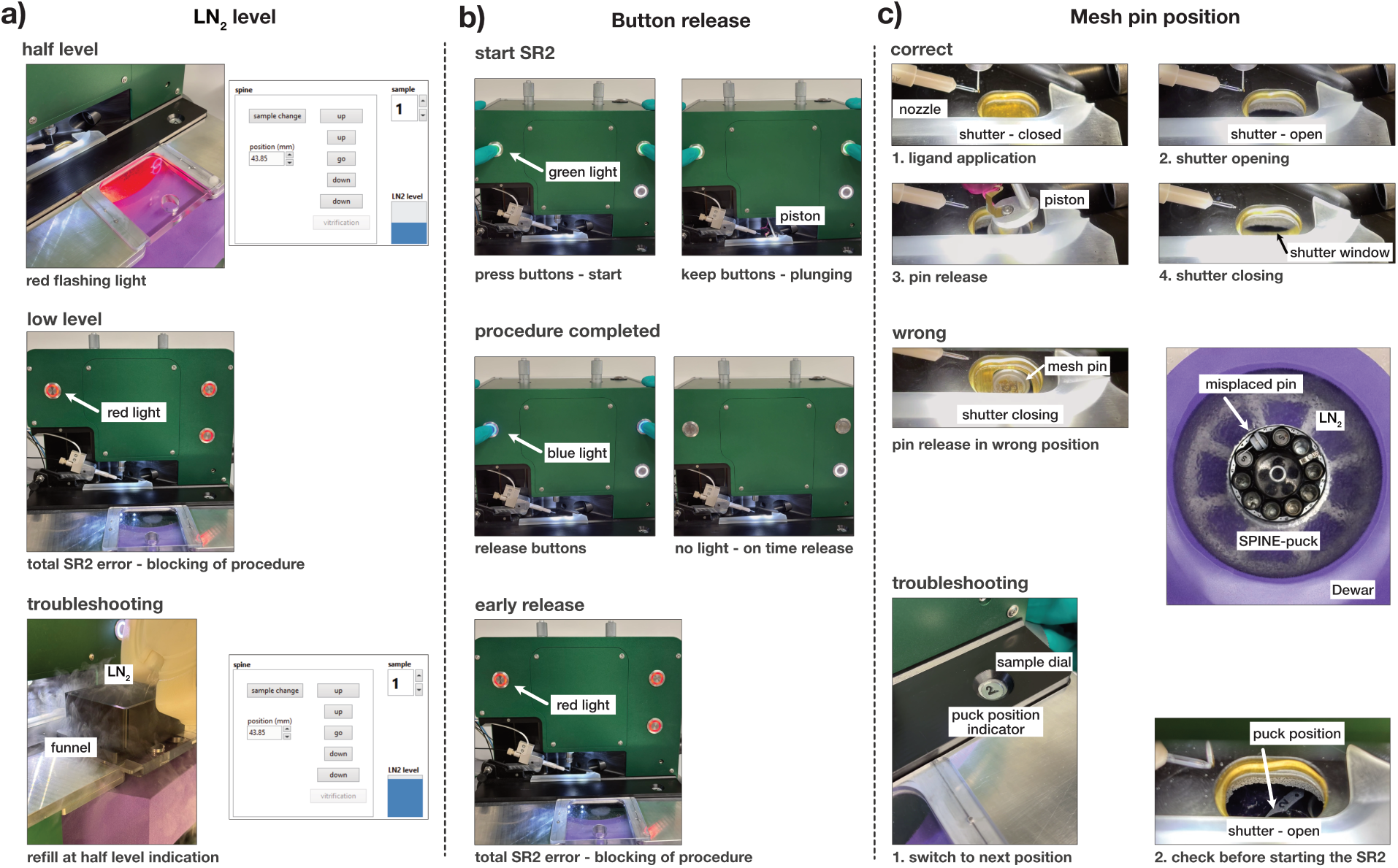
*Spitrobot* troubleshooting. a) Low LN_2_ causes a constant drift in the freezing profile as LN_2_ boils away. Adding to time-delay variation, which particularly effects sub-second time-delays. The *Spitrobot-2* includes an automatic detection for the reduction in LN_2_, when the LN_2_ is low an initial flashing stage warns the experimenter. If the LN_2_ level becomes low enough all buttons turn red at which point the *Spitrobot* will not begin the automated vitrification procedure, until the LN_2_ is replenished. b) Once the trigger buttons have been depressed they must be maintained until the vitrification cycle has completed indicated by the buttons turning blue, this is a deliberate safety feature. Releasing the triggers halts the procedure and the *Spitrobot* has to be reset before further vitrificaton can proceed. A mechanical complication c) upon vitrification can cause the pin to by misplaced either by being trapped in the shutter mechanism or not correctly entering the SPINE puck.

27. Check the liquid nitrogen fill gauge, if it is no-longer at maximum fill using the LN_2_ funnel. If this gauge falls below a certain level while mounting or aligning the next sample it will prevent triggering plunge freezing until the LN_2_ is refilled while the sample is mounted. This should be avoided as LN_2_ boiling off from the funnel can cool the sample despite the stable environment provided by the HFD.

28. Move the Autodrop pipette back a few millimetres with the stages before pulling it back to the retracted position and fixing it with the nozzle pin.

29. A new time-delay setup can now be requested step 17, if the time-delay remains the same, a new *Spitrobot* sample can now be mounted step 20. If the puck is complete, either because its full or no more samples are required, the foam dewar can be removed and the puck stored in a transport dewar for shipping to the beamline. Loading a new puck is outlined in additional protocols in the SI.

### 5.4 Data-collection strategies for cryo-trapped samples

Data-collection for *Spitrobot* samples follows the same procedures already developed for modern MX-beamlines. [60]. Guidelines and protocols on how optimise data collection on modern X-ray source have been extensively addressed elsewhere [61–64]. There are specific considerations for data collections from microcrystals deposited on micromeshes. In this case there three potential data-collection strategies. Option A, crystals are collected as rotation datasets, for samples with ice or extensive residual ligand solution can present difficulties with alignment; Option B, multiple rotation wedges collected on separate crystals and merged together; or Option C the micromesh can be rastered to collect a cryo-SSX dataset.

#### 5.4.1 Rotation datasets from micromeshes

**Option A: Full rotation datasets**

i. Mount the micromesh and align the face perpendicular to the beam, at this stage the crystals are either clearly visible or, as often with microcrystals, maybe difficult to visualise. If crystal positions are clear, align and collect a rotation dataset, otherwise gridscan as outlined below.
ii. Select a small beam-size and raster the mesh, most beamlines have this capacity, and can display the grid-scan as a heat map of diffraction containing images.
iii. For single crystals clearly visible in the heatmap, align the beam to this position and adjust to the beamsize to crystal size, otherwise align to well diffracting areas and sculpt the beamsize as guided by the grid scan.
iv. If there is only moderate depth of the sample compared to the beamsize centre on this position then collect, otherwise perform a second grid-scan at 90 degrees to ascertain the crystal depth on the grid. Use both to align the crystal.

**Option B: A series of wedge dataset collections**

i. Repeat steps (i)-(iii) from Option A
ii. Rather than a full rotation, at each strong diffraction point collect a small rotation (5-30 degrees). These can be scaled and merged into a single dataset using XDS (BLEND) or dials.multiplex [65–67].

#### 5.4.2 Cryo-SSX

**Option C: rastered serial dataset collections**

i. Mount the micromesh and align the face perpendicular to the beam, utilising a microfocus beam if possible or a smaller beam size raster the micromesh
ii. Rotate the mesh 30 degrees and repeat (i), rotate the sample back the other way 60 degrees and repeat (i). This should minimise preferential orientation and increase the total number of lattice orientations sampled.

### 6 Anticipated Results

Automated cryo-trapping experiments will produce at least one puck for control data and then 5 samples per delay times^1^. These can be transferred to unipucks to meet the beamline requirements. Sufficient cryo-protection should produce a clear droplet where crystals can be visualised or aligned via X-ray centring procedures. 2*F_o_*-*F_c_* electron density maps for time-delays should show electron-density for the positive control and this can be tracked as time-delays are shortened. Provided the underlying physical chemistry discussed in the experimental design is amenable to millisecond mixing the presence of ligand should be observed into this regime Figure 7.

**Figure 7:**
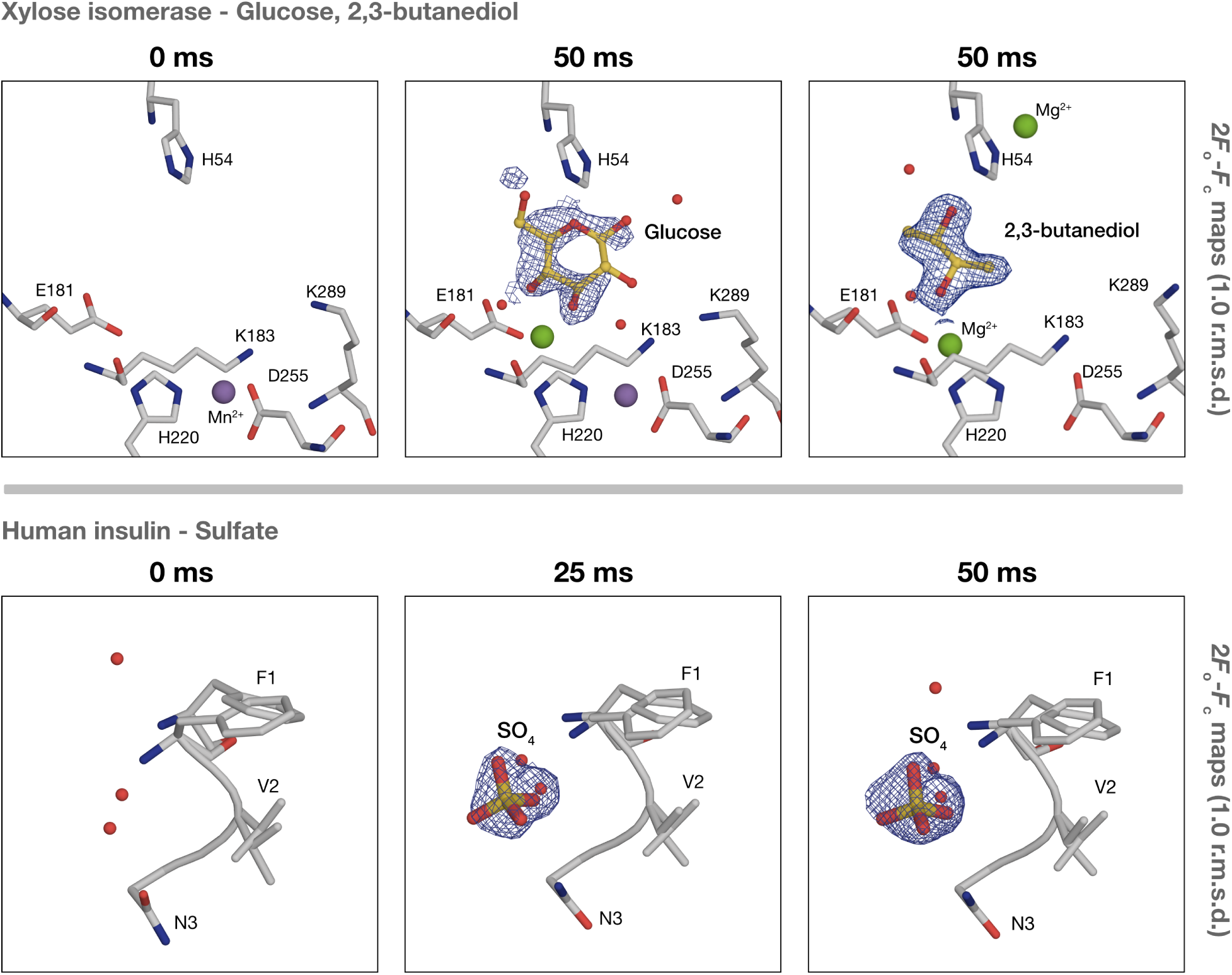
Results from a cryo-trapping experiment on xylose isomerase and human insulin micro-crystals. 2*F_o_*-*F_c_* maps contoured to 1 r.m.s.d of the active site show the presence of the glucose ring after 50 ms (PDB codes: 9R45 (apo state) and 9R47 (50 ms)). Xylose isomerase microcrystals where triggered with 2 M glucose and data was collected via rotation on a micromesh. A separate case shows the 2*F_o_*-*F_c_* map for a control where only the cryo-protectant 2,3-butandiol was present in the buffer (PDB code: 8AWY). The cryo-protectant was shown to bind in the active site, and outcompeted the ligand. This allowed a new cryo-condition to be optimised for subsequent experiments. Human insulin microcrystals were triggered with low pH buffer supplemented with 1 M Na_2_SO_4_ and collected as single crystal rotation data. 2*F_o_*-*F_c_* maps contoured to 1 r.m.s.d of the active site show the presence of the SO_4_ (PDB codes: 9R48 (apo state), 9R49 (25ms), and 9R4A (50 ms)). This figure is adapted from [48, 50].

### 6.1 Data Analysis

Approaches to analyse cryo-trapping data are no different to strategies for time-resolved work in general and therefore we will not cover them in detail here. A standard 2*F_o_*-*F_c_* map should reveal significant intermediates, for comparability a standard *R_free_* set should be maintained. Absolute value END-RAPID maps ensure datasets are on a comparable scale for interrogating changes between time-delays [68]. For earlier delays where ligand occupancy is anticipated to be low, OMIT or POLDER-OMIT (POLDER) maps can delineate the early stages of ligand binding [69]. A straight-forward approach is to calculate all these once a model is reasonably well refined to cross-check consistency between maps.

It is typical to calculate a difference density or *F_o_*-*F_o_* map for photo-triggered time-resolved work. The calculation of *F_o_*-*F_o_* maps has been applied appropriately to cryo-trapping on rhodopsins. This can be useful in mixing experiments, however coordinated waters or ions are often present in the active sites and are therefore subtracted against the ligand. Therefore, *F_o_*-*F_o_* maps have limited utility in interpreting active-site chemistry. Instead they are best for assessing the structural changes to the protein and high-occupancy non-dissociating co-factors present in the apo or ground state that undergo changes during the reaction.

## 7 Conclusions

As the importance in dynamic structural biology grows, the value in studying proteins undergoing catalysis will increase. As such utilising regular and more advanced cryo-trapping approaches to provide additional structural data on protein functional states provides a promising direction for research. The protocol outlines considerations for when and how to apply this methodology, both as a very simple bench-top measurement or utilising automated cryo-trapping as straight forward and relatively robust way to reliably reach time-delays unachievable manually. The *Spitrobot* is an automated cryo-trapping device which has been specifically designed to facilitate these measurements as a standard bench-top device for use alongside other commonly used biochemical tools. This therefore envisages a point when time-resolved crystallography will be a centre piece measurement for integrative structural biology of protein dynamics.

## Acknowledgements

Figures 3 and 5 were created in BioRender. Prester, A. (2026) https://BioRender.com/ybbjyfi

## Funding

The authors gratefully acknowledge the support provided by the Max Planck Society. PM acknowledges support from the Deutsche Forschungsgemeinschaft (DFG) via grant No. 451079909 and from a Joachim Herz Stiftung add-on fellowship. ES acknowledges support by the Federal Ministry of Education and Research, Germany, under grant number 01KI2114. Funded by the European Union. Views and opinions expressed are however those of the author(s) only and do not necessarily reflect those of the European Union or the European Research Council Executive Agency (ERCEA). Neither the European Union nor the granting authority can be held responsible for them.

## 8 Supplementary Information

### 8.1 Crystal mount selection

Provided that a loop can be mounted onto SPINE style magnetic base, it is compatible with the *Spitrobot*. While large macrocrystals can be fished in nylon or Kapton loops, micro-crystals are best loaded onto Micro-mounts, or Micro-Meshes. Standard crystal fishing considerations apply here, such as tailoring the loop size to the crystal. The *Spitrobot* is compatible with regular SPINE mounts large (500 µm) or small (5-40 µm) crystals of different morphologies can be easily mounted as for standard data-collection. Crystals can be fished directly from crystal plates into nylon or Kapton loops, while microcrystals in slurries are typically pipetted directly on Kapton meshes with remaining mother liquor removed via blotting.

### 8.2 Micromeshes

Micromeshes are preferred for mounting microcrystals grown in batch. Slurries can be directly pipetted onto micromeshes and excess mother liquor easily removed via blotting with the mesh preventing crystals being removed by the blotting paper. Hundreds to thousands of crystals can be mounted simultaneously allowing the mesh be rastered for a cryo-SSX strategy, or rotated on individual crystals depending on experimental requirements. As a solid support, meshes can also be glow-discharged to better spread the crystal slurry or ligand solution across the mesh.

### 8.3 Setting up ancillary devices

#### 8.3.1 Setting up the Humidity Flow Device (HFD)

The HFD device provides humidity and temperature control at the sample point and needs to be equilibrated and mounted prior to starting experiments. Most protein crystals survive at 20°C and 95 % humidity which is the assumed default, unless crystals are grown under different conditions (4°C) or where pre-explored environmental conditions are desired [70]. Start the HFD 30 minutes before sample preparation to allow time for equilibration.

1. Using an Allen key unscrew the HFD hose port on the front of the *Spitrobot*, fit the HFD nozzle into the port and position 5-10 cm from the sample position, re-tighten the set screw.
2. Attach a compressed air-line to the back of the HFD. Confirm airflow by adjusting the ’humid air’ and ’dry air’ flow meters to pre-calibrated requirements for temperature and humidity.
3. Fill a 60 mL syringe with deionised water and attach to the HFD water port, fill the waterbath until above the minimum volume as indicated on the side.
4. Using the display set the HFD to ’running’, and navigate to the designated panel to set the ’des’ values to the required temperature and humidity.
5. Place a hand in front of the HFD nozzle to confirm that humidified air is being produced.
6. Wait for around 15 - 30 minutes for the temperature and humidity to equilibrate to the required values, if this seems to be taking longer troubleshoot as in Table 2.

**Table 2:**
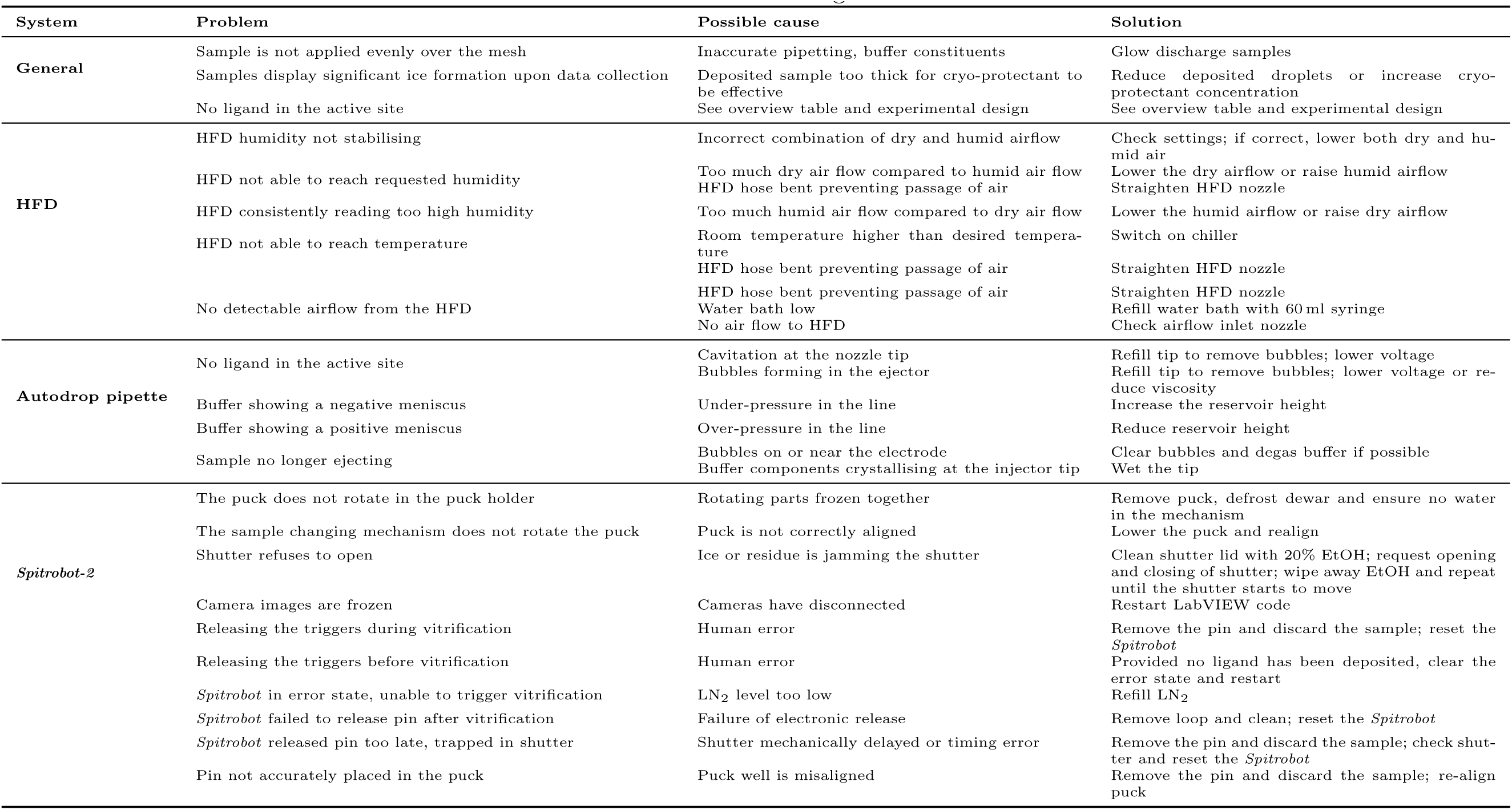
Troubleshooting overview.

#### 8.3.2 Autodrop pipette configuration - Confirm droplet formation

The Autodrop pipette needs to be flushed with the appropriate spitting solution (buffer or ligand), spitting confirmed, and mounted to the device:

1. Fill the Autodrop pipette reservoir with water, flush the line with water to confirm there are no blockages.
2. Attach the Autodrop pipette and flow water through the nozzle to establish a laminar flow.
3. Refill the reservoir with ligand containing buffer (or buffer if performing controls), and repeat 1-2 with the new solution.
4. Adjust the reservoir height to ensure a slight over pressure at the pipette tip.
5. Using the piezo-injector controls, manually trigger a single droplet, visualise this on the water sensitive paper (Figure 4a), if this fails troubleshoot as in Table 2.
6. Once single droplets are established trigger a droplet burst for the desired deposited volume, repeat this 3-4 times to confirm consistency.
7. Leave the droplet nozzle in a humid environment to prevent the tip drying up while setting up the *Spitrobot*.
8. **PAUSE POINT**: If the HFD is already mounted and running, the Autodrop injector can be mounted and left in the humidified stream to prevent buffer evaporation and corresponding crystallisation of buffer components, blocking the injector (Figure 4b).

### 8.4 Optimising the Autodrop pipette for ejection

Crystallisation buffers have different constituents and therefore viscosities and this effects ejection, therefore prior to spitroboting the buffer should be optimised. In-order to form controlled plnl droplets we utilize a piezoelectric injector, similar to other rapid mixing systems [28, 71, 72]. These ejection strategies are sensitive to the buffer viscosity, for example, the spitrobot injector is inconsistent when large molecular weight PEGs are present at over 20% (v/v). As such, the injection parameters must be optimized for each buffer.

1. Fill the Autodrop pipette reservoir with water, flush the line with water to confirm there are no blockages.
2. Attach the Autodrop pipette and flow water through the nozzle to establish a laminar flow.
3. Refill the reservoir with buffer, and repeat 1-2 with the new solution.
4. Start with Autodrop pipette single pulse mode, voltage=160, pulse width=20, pulse delay=9. Around 1000Hz and 200-300 drops. This should deposit around 19 nl.
5. If the sample ejects repeatably then no further work is needed. If no droplet is observed then the flow diagram lays out the iterative optimisation.
6. Once reliable droplet ejection has been established re-clean the lines with water and EtOH.

**Figure 8:**
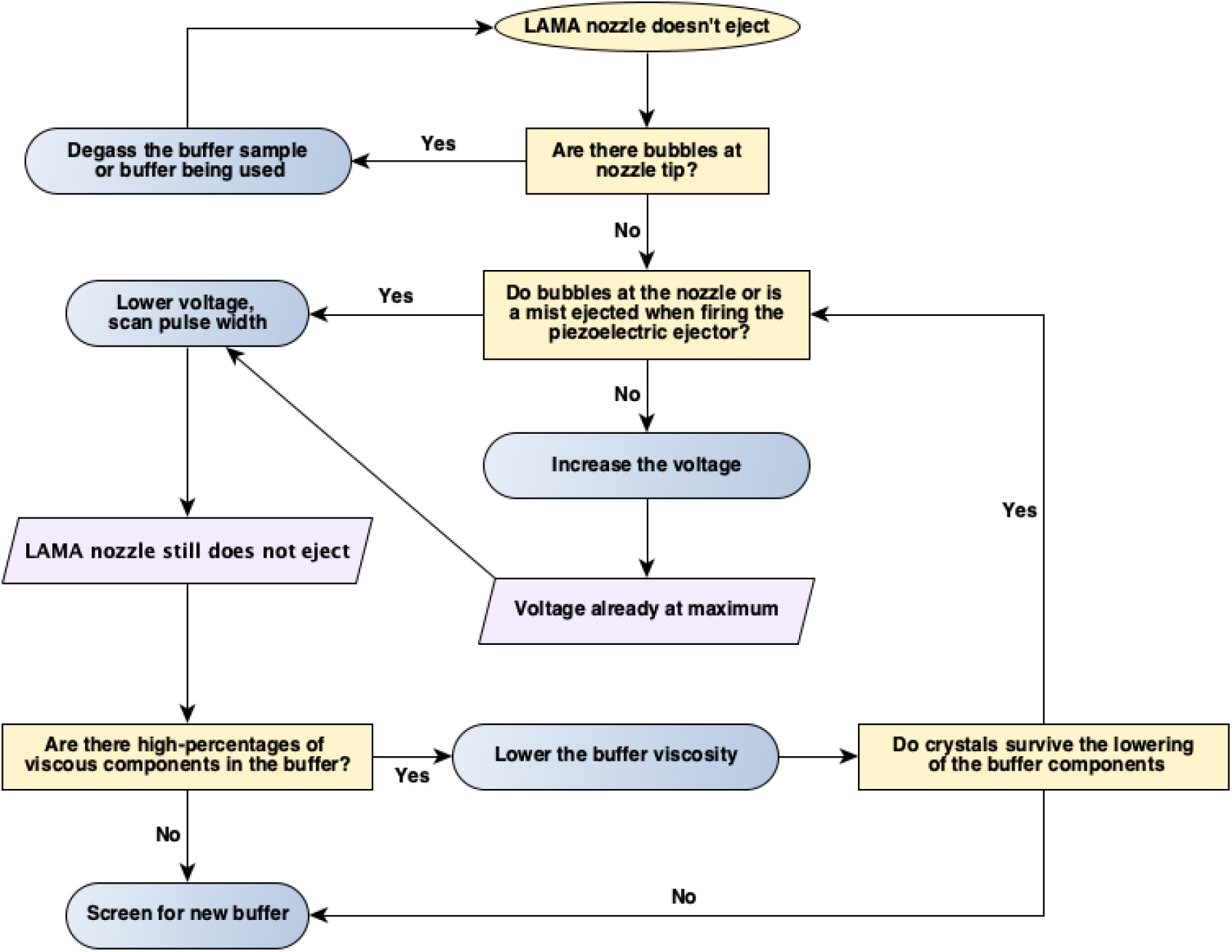
Overview workflow for optimising nozzle ejection. The process is a series of optimisations to deal with buffer viscosity and high solute concentration.

## 9 Materials

## 10 Troubleshooting

## Footnotes

1 As a rule-of-thumb, 5 samples should be prepared per time-point and at least 3 should be collected for cross-comparison.

## Notes

### Competing Interest Statement

The authors have declared no competing interest.

## References

(1) Edman, K.; Nollert, P.; Royant, A.; Belrhali, H.; Pebay-Peyroula, E.; Hajdu, J.; Neutze, R.; Landau, E. M. Nature 1999, 401, 822–826.

(2) Fedorov, R.; Schlichting, I.; Hartmann, E.; Domratcheva, T.; Fuhrmann, M.; Hegemann, P. Biophysical Journal 2003, 84, 2474–2482.

(3) Fink, A. L.; Ahmed, A. I. Nature 1976, 263, 294–297.

(4) Makinen, M. W.; Fink, A. L. Annual Review of Biophysics 1977, 6, 301–343.

(5) Hajdu, J.; Acharya, K.; Stuart, D.; McLaughlin, P.; Barford, D.; Oikonomakos, N.; Klein, H.; Johnson, L. The EMBO Journal 1987, 6, 539–546.

(6) Schlichting, I.; Almo, S. C.; Rapp, G.; Wilson, K.; Petratos, K.; Lentfer, A.; Wittinghofer, A.; Kabsch, W.; Pai, E. F.; Petsko, G. A.; Goody, R. S. Nature 1990, 345, 309–315.

(7) Gouet, P.; Jouve, H.-M.; Williams, P. A.; Andersson, I.; Andreoletti, P.; Nussaume, L.; Hajdu, J. Nature Structural Biology 1996, 3, 951–956.

(8) Käck, H.; Gibson, K. J.; Lindqvist, Y.; Schneider, G. Proceedings of the National Academy of Sciences 1998, 95, 5495–5500.

(9) Schneider, T. R.; Gerhardt, E.; Lee, M.; Liang, P.-H.; Anderson, K. S.; Schlichting, I. Biochemistry 1998, 5394–5406.

(10) Wilmot, C. M.; Hajdu, J.; McPherson, M. J.; Knowles, P. F.; Phillips, S. E. V. Science 1999, 286, 1724–1728.

(11) Wilmot, C. M.; Pearson, A. R. Current Opinion in Chemical Biology 2002, 6, 202–207.

(12) Molina, R.; Stella, S.; Redondo, P.; Gomez, H.; Marcaida, M. J.; Orozco, M.; Prieto, J.; Montoya, G. Nat Struct Mol Biol 2015, 22, 65–72.

(13) Wilmouth, R. C.; Edman, K.; Neutze, R.; Wright, P. A.; Clifton, I. J.; Schneider, T. R.; Schofield, C. J.; Hajdu, J. Nature Structural Biology 2001, 8, 689–694.

(14) Berglund, G. I.; Carlsson, G. H.; Smith, A. T.; Szöke, H.; Henriksen, A.; Hajdu, J. Nature 2002, 417, 463–468.

(15) Pearson, A. R.; Mozzarelli, A.; Rossi, G. L. Current Opinion in Structural Biology 2004, 14, 656–662.

(16) Rasmussen, B. F.; Stock, A. M.; Ringe, D.; Petsko, G. A. Nature 1992, 357, 423–424.

(17) Moffat, K. Chemical Reviews 2001, 101, 1569–1581.

(18) Caramello, N.; Royant, A. Acta Crystallographica Section D: Structural Biology 2024, 80.

(19) Kouyama, T.; Nishikawa, T.; Tokuhisa, H.; Okumura, H. Journal of Molecular Biology 2004, 335, 531–546.

(20) Crosson, S.; Moffat, K. The Plant Cell 2002, 14, 1067–1075.

(21) Hekstra, D. R. Annual review of biophysics 2023, 52, 255–274.

(22) Moffat, K.; Henderson, R. Current Opinion in Structural Biology 1995, 5, 656–663.

(23) Chapman, H. N. Annual review of biochemistry 2019, 88, 35–58.

(24) Wilson, M. A. Annual Review of Biophysics 2022, 51, 79–98.

(25) Gati, C.; Bourenkov, G.; Klinge, M.; Rehders, D.; Stellato, F.; Oberthür, D.; Yefanov, O.; Sommer, B. P.; Mogk, S.; Duszenko, M.; Betzel, C.; Schneider, T. R.; Chapman, H. N.; Redecke, L. IUCrJ 2014, 1, 87–94.

(26) Stellato, F. et al. IUCrJ 2014, 1, 204–212.

(27) Coquelle, N.; Brewster, A. S.; Kapp, U.; Shilova, A.; Weinhausen, B.; Burghammer, M.; Colletier, J. P. Acta Crystallographica Section D: Biological Crystallography 2015, 71, 1184– 1196.

(28) Mehrabi, P.; Schulz, E. C.; Agthe, M.; Horrell, S.; Bourenkov, G.; von Stetten, D.; Leimkohl, J.-P.; Schikora, H.; Schneider, T. R.; Pearson, A. R., et al. Nature methods 2019, 16, 979– 982.

(29) Mehrabi, P.; Müller-Werkmeister, H. M.; Leimkohl, J.-P.; Schikora, H.; Ninkovic, J.; Krivokuca, S.; Andriček, L.; Epp, S. W.; Sherrell, D.; Owen, R. L., et al. Journal of Synchrotron Radiation 2020, 27, 360–370.

(30) Weinert, T.; Skopintsev, P.; James, D.; Dworkowski, F.; Panepucci, E.; Kekilli, D.; Furrer, A.; Brünle, S.; Mous, S.; Ozerov, D.; Nogly, P.; Wang, M.; Standfuss, J. Science 2019, 365, 61–65.

(31) Beyerlein, K. R.; Dierksmeyer, D.; Mariani, V.; Kuhn, M.; Sarrou, I.; Ottaviano, A.; Awel, S.; Knoska, J.; Fuglerud, S.; Jönsson, O., et al. IUCrJ 2017, 4, 769–777.

(32) Prester, A. et al. Communications Chemistry 2024, 7, 1–12.

(33) Evans, G.; Axford, D.; Waterman, D.; Owen, R. L. Crystallography Reviews 2011, 17, Publisher: Taylor & Francis eprint: 10.1080/0889311X.2010.527964, 105–142.

(34) Grünbein, M. L.; Stricker, M.; Nass Kovacs, G.; Kloos, M.; Doak, R. B.; Shoeman, R. L.; Reinstein, J.; Lecler, S.; Haacke, S.; Schlichting, I. Nature Methods 2020, 17, 681–684.

(35) Barends, T. R. M. et al. Nature 2024, 626, 905–911.

(36) Schmidt, M. et al. Advances in Condensed Matter Physics 2013, 2013.

(37) Olmos, J. L. et al. BMC Biology 2018, 16, 1–15.

(38) Pandey, S. et al. IUCrJ 2021, 8, 1–18.

(39) Malla, T. N. et al. Nature Communications 2023, 14, 5507.

(40) Kriminski, S.; Caylor, C.; Nonato, M.; Finkelstein, K.; Thorne, R. Acta Crystallographica Section D: Biological Crystallography 2002, 58, 459–471.

(41) Kriminski, S.; Kazmierczak, M.; Thorne, R. E. Acta Crystallographica Section D 2003, 59, 697–708.

(42) Halle, B. Proceedings of the National Academy of Sciences 2004, 101, 4793–4798.

(43) Indergaard, J. A.; Mahmood, K.; Gabriel, L.; Zhong, G.; Lastovka, A.; McLeod, M. J.; Thorne, R. E. IUCrJ 2025, 12, 372–383.

(44) Warkentin, M.; Berejnov, V.; Husseini, N. S.; Thorne, R. E. Journal of Applied Crystallography 2006, 39, 805–811.

(45) Berejnov, V.; Husseini, N. S.; Alsaied, O. A.; Thorne, R. E. Journal of Applied Crystallography 2006, 39, 244–251.

(46) Ryan, K. P. Scanning microscopy 1992, 6, 8.

(47) Clinger, J. A.; Moreau, D. W.; McLeod, M. J.; Holyoak, T.; Thorne, R. E. IUCrJ 2021, 8, 784–792.

(48) Mehrabi, P.; Sung, S.; von Stetten, D.; Prester, A.; Hatton, C. E.; Kleine-Döpke, S.; Berkes, A.; Gore, G.; Leimkohl, J.-P.; Schikora, H., et al. Nature Communications 2023, 14, 2365.

(49) Kondo, Y., et al. Protein Science 2025, 34, e70104.

(50) Spiliopoulou, M.; Hatton, C. E.; Kollewe, M.; Leimkohl, J.-P.; Schikora, H.; Tellkamp, F.; Mehrabi, P.; Schulz, E. C. Spitrobot-2 advances time-resolved cryo-trapping crystallography to under 25 ms, 2025.

(51) Bourgeois, D.; Royant, A. Current opinion in structural biology 2005, 15, 538–547.

(52) Shoeman, R. L.; Hartmann, E.; Schlicting, I. Nature Protocols 2023, 18, 854–882.

(53) Martin, R. W.; Zilm, K. W. Journal od Magnetic Resonance 2003, 165, 162–174.

(54) Stohrer, C.; Horrell, S.; Meier, S.; Sans, M.; Stetten, D. V.; Hough, M.; Goldman, A.; Monteiro, D. C.; Pearson, A. R. Acta Crystallographica Section D: Structural Biology 2021, 77, 194–204.

(55) Tremlett, C. J.; Stubbs, J.; Stuart, W. S.; Stewart, P. D. S.; West, J.; Orville, A. M.; Tews, I.; Harmer, N. J. IUCrJ 2025, 12, 262–279.

(56) Beale, J. H.; Bolton, R.; Marshall, S. A.; Beale, E. V.; Carr, S. B.; Ebrahim, A.; Moreno-Chicano, T.; Hough, M. A.; Worrall, J. A.; Tews, I.; Owen, R. L. Journal of Applied Crystallography 2019, 52, 1385–1396.

(57) Garman, E. F.; Doublié, S. In *Methods in Enzymology* ; Macromolecular Crystallography, Part C, Vol. 368; Academic Press: 2003, pp 188–216.

(58) Geremia, S.; Campagnolo, M.; Demitri, N.; Johnson, L. N. Structure 2006, 14, 393–400.

(59) Juers, D. H.; Ruffin, J. Journal of Applied Crystallography 2014, 47, 2105–2108.

(60) Cipriani, F. et al. Acta Crystallogr D Biol Crystallogr 2006, 62, 1251–1259.

(61) Garman, E.; Sweet, R. M. In Macromolecular Crystallography Protocols: Volume 2: Structure Determination; Humana Press: Totowa, NJ, 2007, pp 63–93.

(62) Winter, G.; Gildea, R. J.; Paterson, N. G.; Beale, J.; Gerstel, M.; Axford, D.; Vollmar, M.; McAuley, K. E.; Owen, R. L.; Flaig, R.; Ashton, A. W.; Hall, D. R. Acta Crystallogr D Struct Biol 2019, 75, 242–261.

(63) Holton, J. M. J Synchrotron Rad 2009, 16, 133–142.

(64) Popov, A. N.; Bourenkov, G. P. Acta Cryst D 2003, 59, 1145–1153.

(65) Kabsch, W. Acta Cryst D 2010, 66, 125–132.

(66) Foadi, J.; Aller, P.; Alguel, Y.; Cameron, A.; Axford, D.; Owen, R. L.; Armour, W.; Waterman, D. G.; Iwata, S.; Evans, G. Acta Cryst D 2013, 69, 1617–1632.

(67) Gildea, R. J.; Beilsten-Edmands, J.; Axford, D.; Horrell, S.; Aller, P.; Sandy, J.; Sanchez-Weatherby, J.; Owen, C. D.; Lukacik, P.; Strain-Damerell, C.; Owen, R. L.; Walsh, M. A.; Winter, G. Acta Cryst D 2022, 78, 752–769.

(68) Lang, P. T.; Holton, J. M.; Fraser, J. S.; Alber, T. Proceedings of the National Academy of Sciences 2014, 111, 237–242.

(69) Liebschner, D.; Afonine, P. V.; Moriarty, N. W.; Poon, B. K.; Sobolev, O. V.; Terwilliger, T. C.; Adams, P. D. Acta Crystallographica Section D: Structural Biology 2017, 73, 148–157.

(70) Russi, S.; Juers, D. H.; Sanchez-Weatherby, J.; Pellegrini, E.; Mossou, E.; Forsyth, V. T.; Huet, J.; Gobbo, A.; Felisaz, F.; Moya, R.; McSweeney, S. M.; Cusack, S.; Cipriani, F.; Bowler, M. W. *J. Struct.* Biol 2011, 175, 236–243.

(71) Butryn, A.; Simon, P. S.; Aller, P.; Hinchliffe, P.; Massad, R. N.; Leen, G.; Tooke, C. L.; Bogacz, I.; Kim, I.-S.; Bhowmick, A., et al. Nature communications 2021, 12, 4461.

(72) Bar-Even, A.; Noor, E.; Savir, Y.; Liebermeister, W.; Davidi, D.; Tawfik, D. S.; Milo, R. Biochemistry 2011, 50, 4402–4410.

